# Vertical Sleeve Gastrectomy Induces Enteroendocrine Cell Differentiation of Intestinal Stem Cells Through Farnesoid X Receptor Activation

**DOI:** 10.1101/2021.04.22.440705

**Authors:** Ki-Suk Kim, Bailey C. E. Peck, Yu-Han Hung, Kieran Koch-Laskowski, Landon Wood, Priya H. Dedhia, Jason R. Spence, Randy J. Seeley, Praveen Sethupathy, Darleen A. Sandoval

## Abstract

Vertical sleeve gastrectomy (VSG) is one of several bariatric procedures that substantially improves glycemia and energy homeostasis. Increased secretion of multiple gut peptides has been hypothesized to be a critical contributor to VSG’s potent effects to reduce body weight and improve glucose regulation. VSG results in an increase in the number of hormone-secreting enteroendocrine cells (EECs) in the intestinal epithelium, but whether this increase is via proliferation or differentiation of EECs and their subtypes remains unclear. Notably, the beneficial effects of VSG are lost in a mouse model lacking the bile acid nuclear receptor, farnesoid X receptor (FXR). FXR is a nuclear transcription factor that has been shown to regulate intestinal stem cell (ISC) function in cancer models, but whether it plays a role specifically in normal intestinal differentiation remains unknown. Therefore, we hypothesized that the VSG-induced increase in EECs is due to changes in intestinal differentiation driven by an increase in bile acid signaling through FXR. To test this, we performed VSG in mice that express eGFP in ISC/progenitor cells and performed RNAseq on GFP-positive cells sorted from the intestinal epithelia. We also assessed changes in EEC number (marked by GLP-1) in mouse intestinal organoids following treatment with bile acids and/or an FXR antagonist. RNA-seq revealed that FXR is expressed in ISCs and that VSG explicitly alters ISC expression of several genes that regulate intestinal secretory cell development, including EEC differentiation. Mouse intestinal organoids treated with bile acids increased GLP-1-positive cell numbers, whereas a potent FXR antagonist blocked this effect. Taken together, these data indicate that VSG drives ISC fate towards EEC differentiation through FXR signaling.

## Introduction

Vertical sleeve gastrectomy (VSG) is the most frequently performed bariatric surgery in the US^1^. VSG patients lose approximately 81% of excess body mass within 1 year, and approximately 38% of patients with type 2 diabetes mellitus achieve post-operative remission^2^. VSG increases postprandial plasma levels of many gut peptide hormones, such as glucagon-like peptide-1 (GLP-1), gastric inhibitory polypeptide (GIP), peptide YY (PYY), and cholecystokinin (CCK), which all play roles in regulating satiety, lipid handling, energy expenditure, and glucose homeostasis^3^. Importantly, this observation is consistent between clinical and preclinical studies, is seen within 2 days following VSG, and is maintained for at least 5 years after surgery^4–8^. Therefore, the drastic increase in plasma gut peptides with VSG is presumed to be the mechanism for successful VSG outcomes. However, a critical question that remains is what drives the post-operative increases in circulating gut hormones.

A number of studies^9–11^ demonstrated an increase in the number of GLP-1-secreting enteroendocrine cells (EECs) after VSG without any gross morphological changes in the intestinal epithelium. Gut peptides are secreted from EECs, which are scattered individually throughout the intestinal epithelium along with many other intestinal cell types (goblet cells, enterocytes, Paneth cells). The intestinal epithelium is continuously renewed by proliferation and differentiation of crypt base columnar intestinal stem cells (ISCs), which can be isolated and sorted by the marker *Lgr5*^12^. The cell-signaling pathways that regulate ISC proliferation and differentiation have been well-described. First, Notch signaling is required for the renewal of ISCs, and thus the activation of Notch signaling induces cell proliferation^12^. In addition, Notch signaling represses the transcriptional regulator, Atoh1, and thereby promotes ISC differentiation toward absorptive enterocytes. Conversely, the inhibition of Notch signaling activates Atoh1 to the differentiation of secretory lineages progenitor cells such as EEC, goblet cells, and Paneth cells^13^. Therefore, the *Notch* and *Atoh1* gene families play a significant role in ISC maintenance and EEC differentiation. However, whether VSG drives ISC fate toward EECs, and what signals drive this change, is unknown.

We previously found that a bile acid (BA) receptor, farnesoid x receptor (FXR), is necessary for the VSG-induced weight loss and improvements in glucose homeostasis^14^. However, the mechanisms by which FXR mediates the metabolic impact of VSG remained elusive. Like gut peptide hormones, many different BA species are increased several-fold after VSG in mice^15^ and humans^16^. Previous work has demonstrated that the taurine-conjugated bile acid, taurocholic acid (TCA), stimulates intestinal epithelial cell proliferation by activation of epidermal growth factor receptor (EGFR), while a secondary bile acid, DCA, inhibits cell proliferation by activating FXR, suggesting that different BA species act via distinct signaling mechanisms to regulate intestinal epithelial homeostasis ^17^. Moreover, FXR signaling has also been found to regulate ISC proliferation in a colorectal cancer mouse model ^18^. Thus, accumulating data suggest that BA signaling is critical for VSG outcomes and also ISC fate. Hence, we hypothesized that the increased secretion of BAs after VSG drives ISC fate towards EECs through FXR signaling, which then results in a drastic increase of various gut peptides.

## Results

As seen in human bariatric patients^3^ and our previous murine work^19^, VSG results in dramatic reductions in body weight (BW) and fat mass without significantly altering lean mass in diet-induced obese (DIO) mice (Fig. 1A-D). VSG-treated mice initially had a significant reduction in food intake, which was restored to the level of their sham-counterparts within 3-weeks after surgery (Fig. 1E). VSG mice also had improved oral glucose tolerance compared to sham surgery mice (Fig. 1F).

**Fig. 1.**
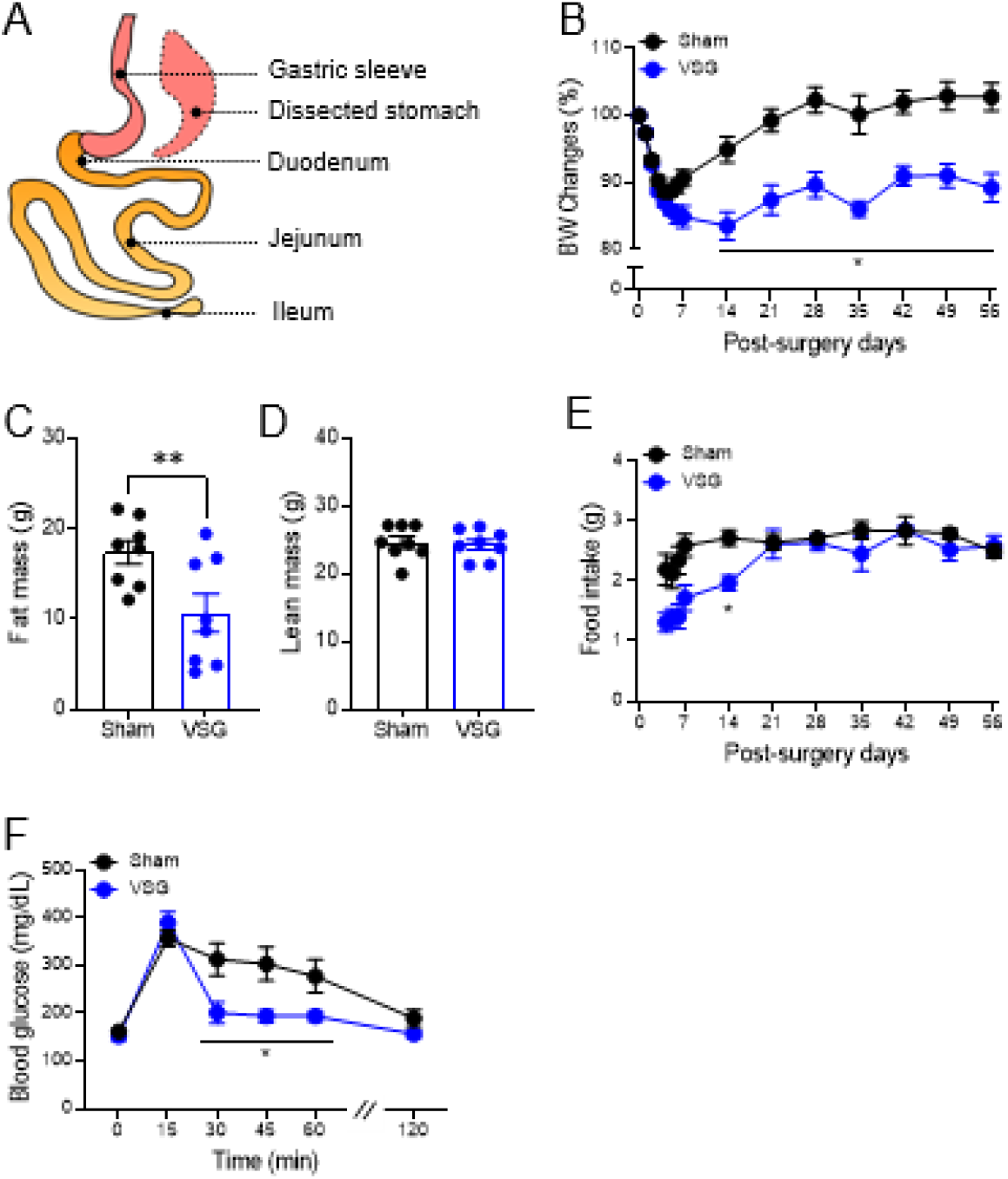
VSG induces metabolic improvements in control male DIO mice. A) A conceptual graphic of VSG procedure. (B-D) BW change from baseline (B), fat mass (C), lean mass (D), and food intake (E) in sham or VSG-treated mice. (F) Oral glucose tolerance (2 g/kg) of sham or VSG-treated mice. (B-C) Body composition was measured at 8 weeks after surgery. (F) Oral glucose tolerance testing was performed at 5-6 weeks after surgery. Mean ± SEM. Statistics, t-test or ANOVA. *P<0.05, **P<0.01, n=8.

Here, we focus on GLP-1 and PYY, as these two anorectic gut peptides are historically suggested to be secreted from the same cells, have demonstrated effects on suppressing feeding, and are increased postprandially by VSG. Consistent with previous studies, plasma total Glp-1 levels were increased 15 min after oral administration of mixed liquid meal in VSG vs. sham mice (Fig. 2A). We also observed that Glp-1-positive (Glp-1^+^) cell numbers were 2-fold higher in VSG vs. sham mice, though this was observed specifically within the jejunal epithelium, with no significant increase in the duodenal or ileal epithelia (Fig. 2B-D). The number of jejunal Glp-1^+^ EECs was significantly associated with plasma levels of total Glp-1 (Fig. 2E). Notably, we did not observe any changes in villi length or crypt depth in any part of the small intestinal sections (Fig. 2F-G), indicating that the increased Glp-1^+^ EEC number is not the result of intestinal proliferation. Plasma levels of Pyy were also increased in VSG-treated mice (Fig. 2H).

**Fig. 2.**
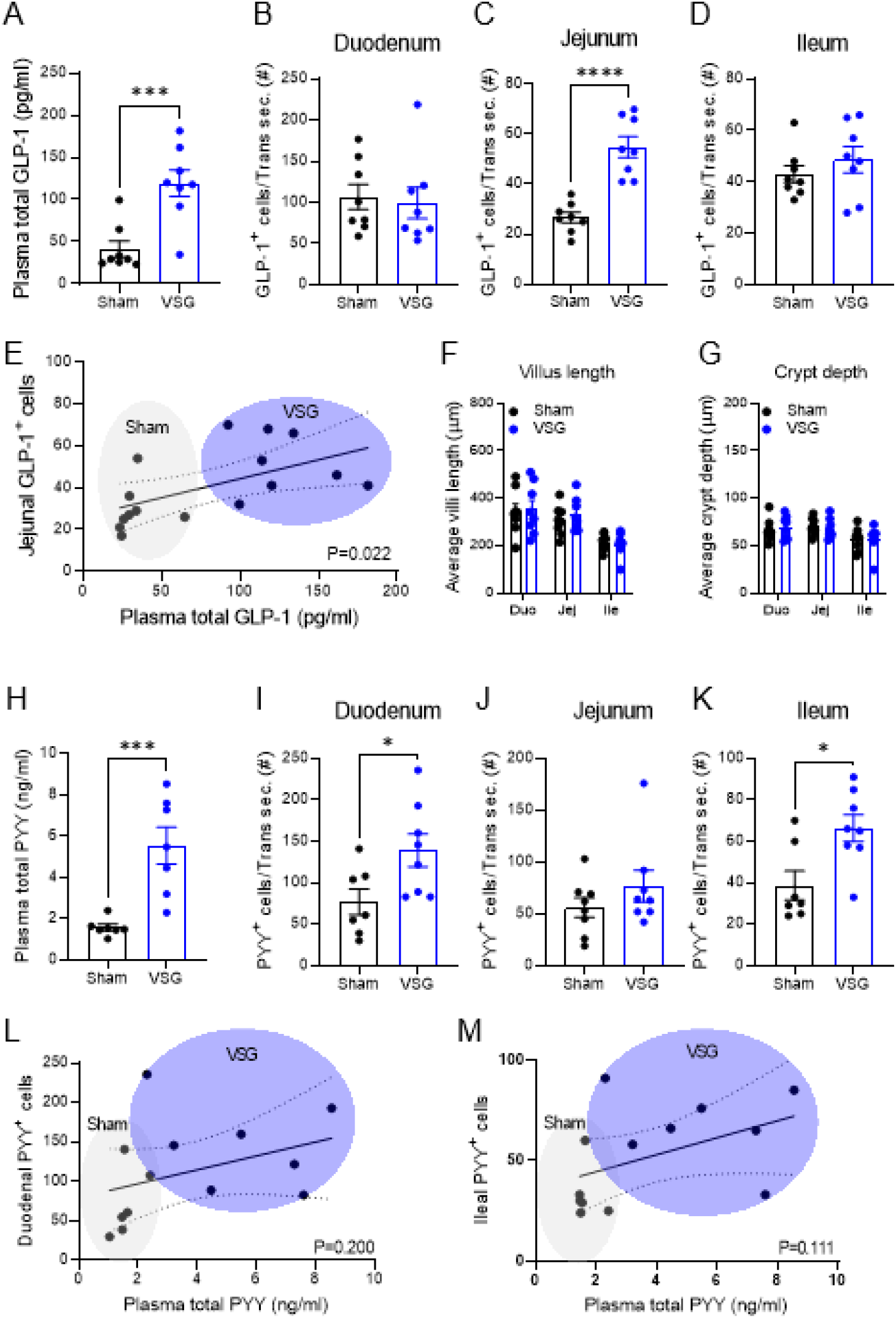
VSG increases plasma levels and EEC number of GLP-1 and PYY. (A) Plasma total GLP-1 levels 15 min after a mixed liquid-meal in sham or VSG-treated mice. (B-D) GLP-1+ EEC number of intestinal transverse sections (Trans sec.) of the duodenum (B), jejunum (C), or ileum (D). (E) Linear regression analysis of the jejunal GLP-1+ cell number vs. plasma total GLP-1 level. (F-G) The villi length (F) and crypt depth (G) of the intestinal segments. (H) Plasma total PYY levels at 15 min after a liquid mixed-meal administration in sham or VSG-treated mice. (I-K) PYY+ EEC number of the intestinal transverse sections of the duodenum (I), jejunum (J), or ileum (K). (L-M) Linear regression analysis of the duodenal (L) or ileal (M) PYY+ cell number vs. plasma total PYY level. Mean ± SEM. Statistics, t-test or ANOVA. *P<0.05, ***P<0.0001, n=8.

Intriguingly, we observed Pyy^+^ EEC number was significantly increased in the VSG-treated duodenal and ileal epithelia, but not in the jejunal epithelium (Fig. 2I-K), demonstrating a regional difference in VSG-induced changes within subtypes of EEC. The number of duodenal or ileal Pyy^+^ EEC did not significantly correlate with the plasma levels of total Pyy (Fig. 2L-M), although there was a trend for a significant correlation when the duodenal and ileal cell numbers were pooled (p=0.08). These data suggest that there are intestinal adaptations to VSG that drive increases in postprandial GLP-1 levels.

Given the high turnover rate of cells within the intestinal epithelium, cellular proliferation and differentiation is a highly regulated process. To determine whether the increase in EECs after VSG was due to alterations in differentiation, we performed VSG on a genetically-modified mouse model expressing eGFP in its *Lgr5* gene (Fig. S1A-B), a specific ISC/progenitor marker gene^12^. As expected, the VSG-treated Lgr5^eGFP^ mice demonstrated reductions in BW and fat mass, normal lean mass, and improvements in oral glucose tolerance compared to their sham-counterparts (Fig. S1C-F). We then performed RNA-seq on the GFP^+^ ISCs sorted from the duodenal, jejunal, and ileal epithelia. We pre-sorted immune cells (CD45 positive), endothelial cells (CD31 positive), apoptotic (Annexin5 positive), and dead cells (stained with Sytox blue) (Fig. S2A) and saw no effect of surgery on the abundance of these cell populations. We did observe that the proportion of GFP^+^ ISCs was decreased in the duodenum and jejunum, but not ileum, in VSG-treated mice (Fig. S2B). Principal component analysis (PCA) of the transcriptomic profiles stratified sham and VSG-treated mice across all segments (Fig. S3A-C), demonstrating the differential impact of VSG on the various sections of the small intestine. By performing differential expression analysis (P < 0.05, adjusted P <0.2, log_2_ fold change > 0.5, normalized count > 50 by DESeq2), we found that VSG altered 467 genes (259 up; 208 down) in the duodenum, 510 genes (348 up; 162 down) in the jejunum, and 955 genes (600 up; 355 down) in the ileum (Fig. S3D-E). Of note, some of the VSG-increased genes are region-specific (Fig. S3G-H). Consistent with our observation that VSG promotes allocation of EECs in the small intestine, we found that GFP^+^ ISCs, particularly from duodenal and jejunal regions, exhibit elevated expression patterns of genes that mark and/or regulate secretory lineage differentiation after VSG (Fig. 3). These include genes that encode proteins involved in secretory progenitor cell fate control (*Atoh1*^12^ and its target genes *Spdef, Dll3*, and *Dll4*) and EEC differentiation (*Pax4*^20^, and *Pax6*^21^), as well as gut peptides (*Gcg, Pyy, Cck, Gip*, and *Fgf15*). In contrast, enterocyte marker genes (*Sis, Alpi, Enpp7*, and *Apoc3*) were decreased in GFP^+^ ISCs of duodenal and jejunal samples from VSG mice (Fig. 3). Of note, several genes involved in Notch signaling, a pathway critical for priming ISCs towards the absorptive lineage^13^, were remarkably decreased after VSG, especially in the jejunum (Fig. 3).

**Fig. 3.**
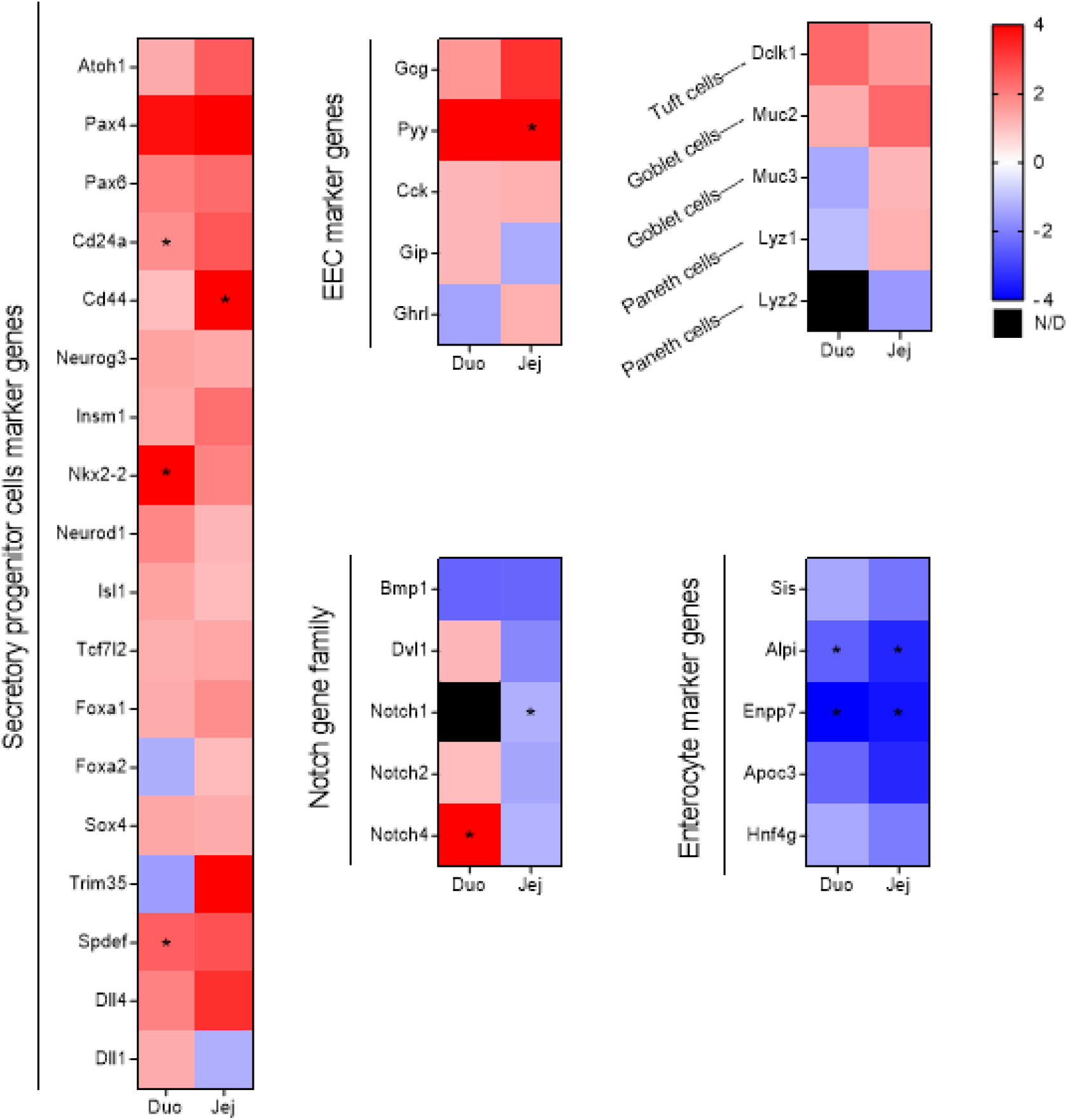
Heatmap analysis of the RNAseq results of the sham or VSG-treated Lgr5-eGFP^+^ cells in duodenum and jejunum. Color denotes variation across stages (normalized by DESeq2). Sham (n=2) and VSG (n=4) were used for RNAseq. Statistical significance in VSG vs. sham was denoted as asterisk when adjusted P value is less than 0.2.

We validated the RNA-seq data with qPCR and confirmed that *Gcg* (GLP-1 encoding gene), but not *Pyy*, was significantly increased in the jejunal ISCs of VSG-treated vs. sham mice (Fig. S4). Though we observed an upregulation of the classic marker of goblet cells (*Muc2*) across all three regional segments (Fig. 3), this change did not lead to the corresponding increase in the protein expression (Fig. S5). These data suggest that VSG induces ISC fate toward EEC differentiation and that there are critical regional differences among ISCs in the duodenum, jejunum, and ileum in the regulation of GLP-1^+^ and PYY^+^ cell differentiation.

To understand whether the signals that are driving differentiation come from the intestinal lumen (which includes ingested nutrients, bile acids, and other digestive enzymes) or the serum (including circulating hormones, cytokines/chemokines, and bile acids), we treated C57BL/6J mouse-derived enteroids with intestinal chyme or sera from sham or VSG-treated rats^23^. Treatment of enteroids with chyme from VSG-treated rats (0.1% vol/vol of the vehicle) significantly increased the enteroid area (Fig. 4A) and the number of GLP-1^+^ cells per enteroid (Fig. 4B) compared to the vehicle-treated control. At the same time, chyme administration did not change the GLP-1^+^ cell density (i.e., GLP-1^+^ cell number per enteroid area) (Fig. 4C). Treatment of enteroids with sera from the sham-treated rats (20% vol/vol of the vehicle) significantly increased enteroid growth, while the sera from the VSG-treated rats did not alter enteroid size (Fig. 4D). Intriguingly, treatment of enteroids with sera from the sham-treated rats did not change the number of GLP-1^+^ cells per enteroids (Fig. 4E), but decreased the GLP-1^+^ cell density, which is restored with sera from the VSG-treated rats (Fig. 4F). These data suggest that both luminal and circulating signals regulate ISC homeostasis.

**Fig. 4.**
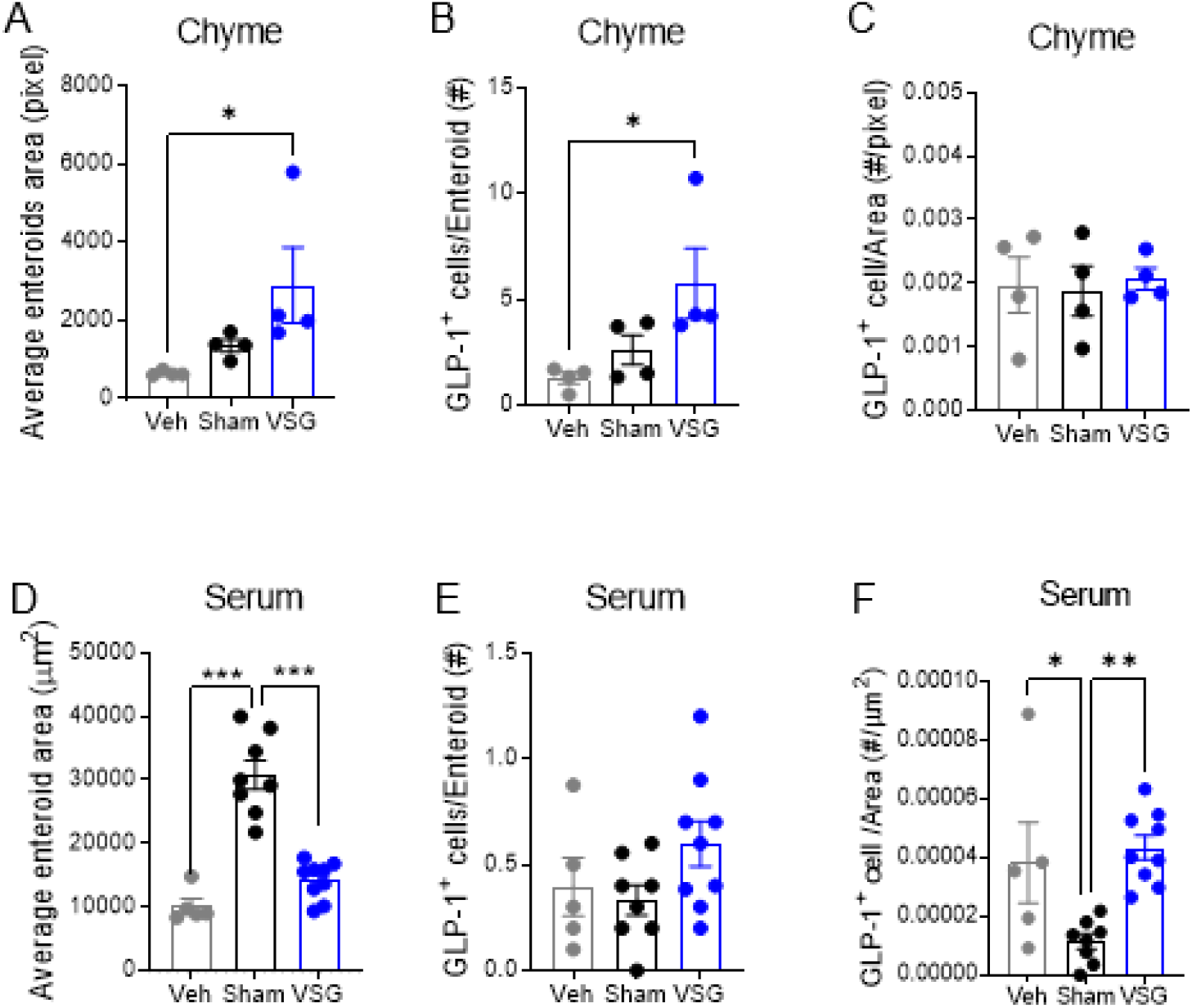
Luminal and serosal factors induce enteroids growth and increase GLP-1+ EEC number in mouse enteroids. (A-C) Chyme from sham or VSG-treated rats was applied to mouse duodenal-derived enteroids for 48 h. The average enteroid area (A), GLP-1+ cell number per enteroid (B), and GLP-1+ cell number per enteroid area (GLP-1+ cell density; C) were measured. (D-I) Sera from sham or VSG-treated rats was applied to mouse duodenal-derived enteroids (20% of the vehicle, vol/vol) for 12 h (D-F). The average enteroid area (D, G), GLP-1+ cell number per enteroid (E, H), and GLP-1+ cell number per enteroid area (F, I) were measured. (A-C) n=4 wells/group, each well contains 10-15 mouse duodenal enteroids. Intestinal chyme was pooled from 3-sham vs. 3-VSG-treated rats and was diluted at 0.1% of the vehicle (vol/vol). (D-I) n=5-10 wells/group, each well contains 10-mouse duodenal enteroids. The sera was pooled from 3-sham and 3-VSG-treated rats and was diluted at 20% of the vehicle (vol/vol). Mean ± SEM. Statistics, ANOVA. *P<0.05, **P<0.01, ***P<0.0001.

BAs are found in both intestinal chyme and serum and are significantly increased after VSG^3^. Therefore, we treated mouse duodenal-derived enteroids with various primary BAs, including cholic acid (CA), chenodeoxycholic acid (CDCA), and muricholic acid (MCA); a secondary BA, DCA; and a tertiary BA, TUD, and examined enteroid growth and changes in GLP-1^+^ cell number. We observed that CA dose-dependently increased GLP-1^+^ EEC number and density, while it did not affect duodenal enteroid area (Fig. 5A-C). Additionally, CA treatment dose-dependently increased the proportion of the enteroids containing GLP-1^+^ EECs (Fig. S6A). CA treatment had similar effects when administered to mouse jejunum-derived enteroids (Fig. S6B). Interestingly, a potent GLP-1 receptor antagonist, exendin 9-39 (Ex9), blocked the effect of CA treatment in increasing GLP-1^+^ EECs number and density (Fig. S7), suggesting that GLP-1 receptor signaling is involved in BA-induced EEC differentiation. Another primary BA, CDCA, increased GLP-1^+^ EEC number and density at a lower dose (1 µM) compared to CA (20 µM), while it did not affect the enteroid area (Fig. 5D-F). However, CDCA treatment did not show the same dose-dependency, nor did it show greater efficacy than the CA treatment (approximately 2-fold increase with CDCA vs. 3-fold with CA). MCA, another major primary BA in mice, dose-dependently increased GLP-1^+^ EEC number and enteroid growth simultaneously, and thus it did not change GLP-1^+^ EEC density (Fig. 5G-I and Fig. S8). DCA and TUD, a secondary and tertiary BA, respectively, did not affect GLP-1^+^ number or density, nor did these BAs affect enteroid area (Fig. 5M-O). These data suggest that different BA species have differential roles in regulating ISC fate and that CA is particularly potent at driving increases in GLP-1^+^ cells.

**Fig. 5.**
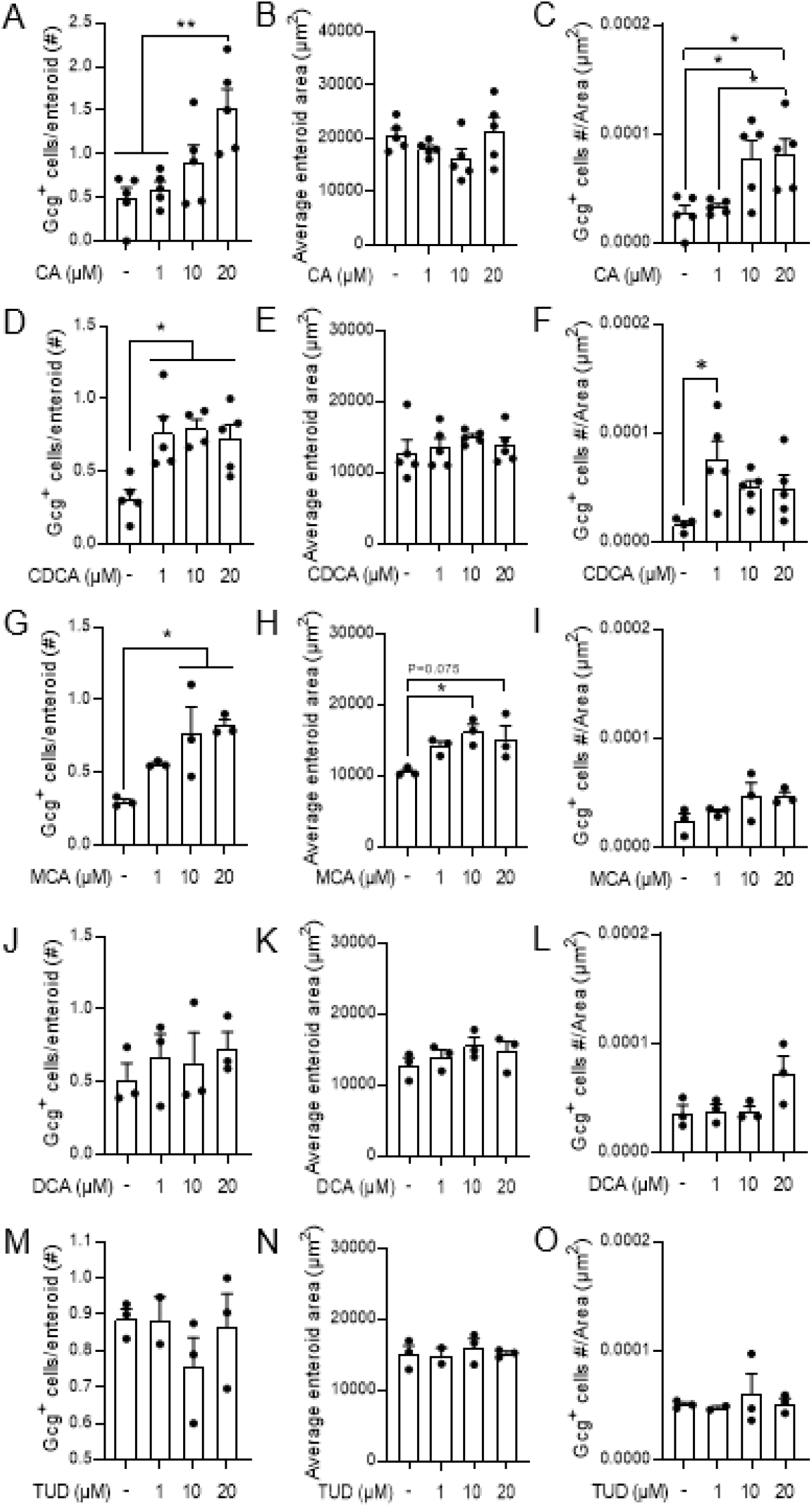
Specific BA treatment increased the growth and GLP-1+ EEC number in the Gcg-tomato (GcgCreERT2 x Rosa26 tdTomato) mouse-derived enteroids. Mouse duodenal enteroids were treated with cholic acid (CA; A-C), chenocholic acid (CDCA; D-F), beta-muricholic acid (MCA; G-I), deoxycholic acid (DCA; J-L), or tauroursodeoxycholic acid (TUD; M-O) for 24 h. The average enteroid area (A, D, G, J, M), GLP-1+ cell number per enteroid (B, E, H, K, N), and GLP-1+ cell number per enteroid area (GLP-1+ cell density; C, F, I, L, O) were measured. n=3-5 wells/group, each well contains 20 enteroids. Mean ± SEM. Statistics, ANOVA. *P<0.05, **P<0.01.

Notably, we found that a number of BA receptor mRNA transcripts were highly expressed in the *Lgr5*+ ISC/progenitor cells throughout the small intestine (Table 1). These BA receptor genes include farnesoid x receptor (*Fxr*), epithelial growth factor receptor (*Egfr)*, pregnane x receptor (*Pxr*), liver X receptor (*Lxr*), vitamin D receptor (*Vdr*), constitutive androstane receptor (*Car*), and retinoid x receptor (*Rxr*) (Table 1), indicating that BAs could regulate ISC fate via a variety of receptors. Because our previous work demonstrated that FXR is necessary for metabolic improvements with VSG, we next tested whether BAs increase GLP-1^+^ EEC number in an FXR-dependent manner. To do this, we administered CA with a potent FXR antagonist, Z-guggulsterone, to mouse-derived enteroids. While the FXR antagonist did not independently influence enteroid area, the effect of CA treatment in increasing GLP-1^+^ EEC number and density was completely blocked (Fig. 6A-D), indicating that FXR activation is necessary for BA-stimulated increases in GLP-1^+^ EEC differentiation.

**Table 1.** Average counts of RNA transcript in FACS-sorted mouse Lgr5^+^ cells (normalized using DESeq2).

| Gene | Duodenum | Jejunum | Ileum |
| --- | --- | --- | --- |
| *Bile acid receptors* |  |  |  |
| *Nr1h4 (Fxr)* | 1357.76 | 2295.54 | 6595.87 |
| *Egfr* | 995.35 | 1317.64 | 322.97 |
| *Gpbar1 (Tgr5)* | N/D | N/D | N/D |
| *Nr1h3 (Pxr)* | 1856.53 | 1760.60 | 2915.18 |
| *Nr1i2(Lxr)* | 4248.62 | 3575.63 | 4553.97 |
| *Vdr* | 18843.76 | 16228.03 | 12987.74 |
| *Nr1i3(Car)* | 2484.43 | 2817.68 | 964.95 |
| *Rxra (Rxr)* | 3578.49 | 4938.92 | 8997.61 |
| *Nr1h4 (Fxr)* | 1357.76 | 2295.54 | 6595.87 |
| *Bile acid signaling receptors* |  |  |  |
| *Fgfr1* | 96.43 | 18.97 | 79.68 |
| *Fgfr2* | 874.66 | 930.76 | 887.38 |
| *Fgfr3* | 1114.45 | 1859.93 | 1193.78 |
| *Fgfr4* | 734.26 | 490.88 | 168.01 |
| *Glp1r* | N/D | N/D | N/D |

**Fig. 6.**
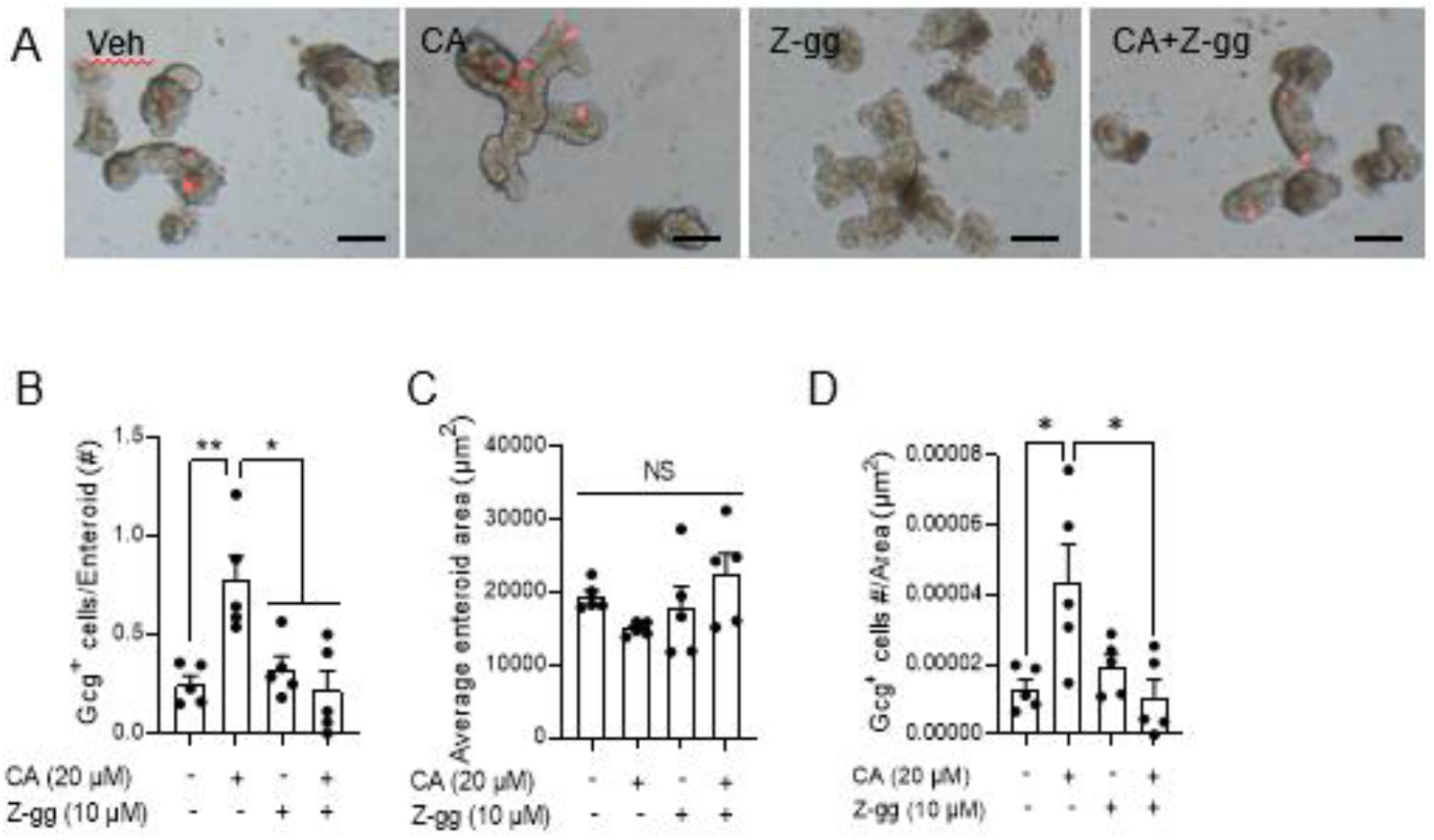
Cholic acid (CA) treatment increases GLP-1+ EEC number in a FXR-dependent manner. (A-D) CA or a potent FXR antagonist, Z-guggulsterone (Z-gg) was administered to Gcg-tomato mouse-derived enteroids for 48 h, and the average enteroid area (B), GLP-1+ cell number per enteroid (C), and GLP-1+ cell number per enteroid area (GLP-1+ cell density; D) were measured. Bar = 100 µm. n=5 wells/group, each well contains 20 enteroids. Mean ± SEM. Statistics, ANOVA. *P<0.05, **P<0.01.

## Discussion

The dramatic increase in postprandial gut peptide levels after VSG is commonly observed in rodent VSG models and in patients who undergo bariatric surgery^3^. These gut peptides have roles in regulating various aspects of energy homeostasis and blood glucose^24^. Despite the fact that the beneficial effects of VSG depend on FXR^14^, the target organ for this BA signaling effect remains elusive. Fundamental questions remain regarding what signal drives the increase in gut peptide levels after VSG and whether this phenomenon is linked to BA-FXR signaling. Here, we demonstrate that VSG increases GLP-1^+^ and PYY^+^ EEC numbers and shifts the gene expression profiles of Lgr5^+^ ISC/progenitor cells toward EEC differentiation. We also demonstrate in enteroids that BA signaling through FXR drives an increase in GLP-1^+^ EEC.

Our group previously demonstrated that gastric emptying rate (GER) is increased after VSG^25^. Many have hypothesized that increases in GER raise gut hormone levels through the early exposure of the proximal (where the majority of GIP and CCK-secreting cells are located) and distal intestinal epithelium (where it is believed the majority of GLP-1- and PYY-secreting cells are located)^26^ to ingested nutrients. However, we found that the same dose and rate of infusion of dextrose directly into the duodenum still causes a significantly greater GLP-1 levels in VSG vs. sham animals ^25^. These data suggest that there are intestinal adaptations that drive the increase in postprandial gut peptide secretion. Our current data suggest that the increased drive towards differentiation of EECs is one such adaptation.

Our murine VSG data demonstrate an increased number of EECs, consistent with some previous studies in rodents^9–11^ and patients^27^. However, one study did not observe this increase in EEC number in a rat model of VSG^28^. The cause of this discrepancy is unclear. While other rodent VSG studies maintained animals on the same pre-surgical high-fat diet (HFD) after surgery, Mumphrey et al. gave their animals a choice between regular chow and HFD postoperatively^28^. HFD and long-chain fatty acids stimulate ISC proliferation^29^, and obesity^27^ or chronic HFD consumption inhibits EEC differentiation and thus reduces EEC number and density^30^. Because VSG^31^ (and another bariatric surgery, Roux en Y gastric bypass^32^) shifts food choice away from fat-rich foods, the reduction in HFD consumption could affect ISC fate. Our data demonstrate that sera from sham rats (which probably contains abundant long-chain fatty acids and other sources of fat) increased enteroid area while it decreased GLP-1^+^ cell density (see Fig.4 D-F). Conversely, the enteroids administered sera from animals that had VSG but were fed HFD had an enteroid area and GLP-1^+^ cell density that was similar to the control (see Fig.4 D-F). Surprisingly, the effect of obesity on ISC fate is reverted by VSG in humans^27^. These data suggest that diet and surgery regulate the composition of circulating factors, which, in turn, regulates ISC fate.

Specific BA subtypes are increased several-fold in sera and jejunum of VSG-treated mice^15^ and in patients that undergo VSG^16^. A growing body of evidence reveals that BAs are not only lipid emulsifiers but are also key signaling molecules with multiple target organs. In particular, studies have demonstrated that BAs regulate ISC fate through FXR and EGFR. An FXR agonist^18^ or specific BAs that activate FXR (e.g., TCA)^17^ inhibited intestinal epithelial proliferation, whereas other BAs that inhibit FXR signaling (T-βMCA and DCA) increased intestinal proliferation^18^, likely through an EGFR-dependent manner^17^. None of these studies evaluated the role of BA receptors in specifically regulating EEC differentiation. Here, we demonstrated that a primary BA, CA, increased GLP-1^+^ EECs in an FXR-dependent manner. Altogether, our data suggest that FXR activation activates EEC differentiation pathways while inhibiting ISC proliferation.

A recent study has demonstrated that the BA subtype, lithocholic acid (LCA) and a GPBAR1 (a.k.a. TGR5) agonist increased GLP-1^+^ cell numbers in mouse-derived enteroids in a GLP-1 receptor- and a serotonin 5-hydroxytryptamine receptor 4 (5-HT4) dependent manner. Conversely, our study demonstrated that the CA-induced increase in GLP-1^+^ cells was blocked by antagonizing the GLP-1 receptor; however, we did not find transcripts for GPBAR1 or GLP-1 receptor within ISCs (see Table 1). Therefore, it is possible that other intestinal epithelial cells that express GLP-1 receptor indirectly regulate EEC differentiation of ISCs. The 5-HT4 gene (*Htr4*) is expressed in ISCs in the duodenum and jejunum (though not in ileal ISCs), but the numbers of the transcripts were very low (152 and 29 transcripts, respectively). Still, it is possible that some BAs directly target ISCs to increase GLP-1^+^ cell number through the 5-HT4 receptor, or they indirectly affect ISC fate through the enterochromaffin cells (that secrete serotonin).

In this study, we demonstrated that VSG changes ISC fate toward EEC differentiation. This change of ISC fate accompanies the upregulation of genes involved in EEC differentiation, the downregulation of genes within the Notch family (the signaling pathway that regulates ISC proliferation and enterocyte differentiation)^13,33^, and the downregulation of enterocyte marker genes. However, there are many different types of EECs, and interestingly, we saw a regional difference between changes in the number of GLP-1^+^ and PYY^+^ cells. The reason for this is unclear. Our RNA-seq analysis reveals that *Foxa2* (a late EEC differentiation marker^22,30^) and *Trim35* (an L-cell marker^22^) were downregulated, and *Dll1* (an early EEC differentiation marker^22^ as well as a Notch ligand^33^) was upregulated within the duodenal vs. jejunal ISCs. Also, the expression of Notch family genes, *Notch1, Notch2*, and *Notch4*, were downregulated by VSG in the jejunum but not the duodenum. Whether these genes are responsible for the differentially expressed EECs across differing regions of the intestine requires further investigation.

Despite the fact that CA has a similar efficacy in increasing GLP-1^+^ cell number in both duodenal- and jejunal-derived enteroids, we only observed an increase in GLP-1^+^ cell number in the jejunum of VSG-treated mice. Previous work has shown that the levels of unconjugated BAs, especially of primary BAs (CA and MCA), drastically increase in the jejunum of VSG-treated mice compared to pair-fed sham-treated mice, and this phenomenon is not observed in the ileum (unclear about the duodenum)^34^. Therefore, it is possible that the difference in BA levels between intestinal regions contributed to the difference in GLP-1^+^ cell number. Why specific BA levels are increased in distinct regions of the intestine after VSG, and whether this change is related to the level of EEC differentiation, requires further investigation.

Furthermore, the increase in jejunal GLP-1^+^ cell number after VSG might have clinical implications. A recent study revealed that jejunal GLP-1^+^ cell number is decreased in obese patients (even more so in obese patients with T2DM) compared to non-obese individuals^35^. Jejunal EECs isolated from patients with T2DM also demonstrate downregulation in genes involved in EEC differentiation (including *PAX4* and *FOXA2*) and in the *GCG* gene itself^35^, indicating that jejunal GLP-1^+^ cell differentiation and number are negatively impacted by both obesity and T2DM.

Many studies, including our previous studies, revealed that GLP-1 or PYY, in and of themselves, are not necessary for the metabolic effects of VSG^19,36,37^. However, many genetic mouse models that target one gut peptide have a compensatory increase in the secretion of other gut peptides (e.g., Gcg-KO mice have a compensatory increase in PYY and GIP levels^19^). Meanwhile, exogenous administration of a combination of multiple gut-peptides (GLP-1, PYY, and oxyntomodulin)^38^ or combined agonists that target multiple gut peptides^39^ have synergistic effects on weight loss and improvements in glycemia. Thus, it may be that surgery-induced increases in multiple gut peptides are necessary for the complete metabolic success of VSG. In addition, the multi-targeting effects of bariatric surgery may play a role in orchestrating the metabolic improvements after VSG.

In conclusion, VSG increases postprandial plasma gut peptide levels, and our data indicate that this is the result of increased number of EECs. Furthermore, we demonstrated that the increase in EEC is driven by VSG-induced changes ISC fate toward EEC differentiation in a FXR-dependent manner.

## Materials and Methods

### Reagents

CA (cat# C1129), CDCA (cat# C9377), MCA (cat# SML2372), DCA (D2510), TUD (cat# 580549), 4-hydroxy tamoxifen Insolution™ (cat# 5082250001), and Z-gg (cat# 370690) were purchased from Millipore Sigma (Burlington, MA, USA). Ex9 was purchased from Bachem (cat# 4017799), Torrance, CA, USA).

### Antibodies

The following antibodies were used in this study: Rabbit anti-GLP-1 (Peninsula Laboratories, # T-4363); Rabbit anti-PYY (Abcam, #ab22663); Rabbit anti-Muc2 (abcam, #ab97386); Rat anti-CD45-APC (Biolegend, #103111); Rat anti-CD31-APC (Biolegend, #102409); APC Annexin V apoptosis detection kit with 7-AAD (Biolegend, #640930); Sytox™ Blue Dead Cell Stain for flow cytometry (ThermoFisher, #S34857); Donkey anti-rabbit conjugated with Alexa 488 (ThermoFisher, #A21206); Goat anti-rabbit IgG conjugated with Cy3 (ThermoFisher, #A10520).

### qPCR primers (or taqman #)

Reverse transcription of mRNA was performed with the High capacity RNA to cDNA kit (Life Technologies, Grand Island, NY). TaqMan assays and TaqMan Gene Expression Master Mix (Life Technologies) were used per the manufacturer’s protocol on a BioRad CFX96 Touch Real Time PCR Detection System (Bio-Rad Laboratories, Richmond, CA). Reactions were performed in triplicate using *B2M* as the normalizer.

#### TaqMan probes

- Gcg Assay ID: Mm01269055_m1
- Pyy Assay ID: Mm00520716_g1

### Animals

All animals were single-housed under a 12-hour light/dark cycle with ad libitum access to water and standard chow (Envigo Teklad; cat# 7012) or 60% HFD (Research Diet; cat# D12492). The animal room was maintained at 25°C with 50%-60% humidity. All studies were performed using animals 8-16 weeks of age, and all animals were euthanized with CO2 inhalation.

Male control mice (VilCre) were maintained on 60% HFD for 6 weeks to induce obesity, were matched for body weight and body fat, and then received sham or VSG surgery as described previously^19^. Briefly, a small laparotomy incision was made in the abdominal wall of the anesthetized mice, and the lateral 80% of the stomach along the greater curvature was excised using an ETS 35-mm staple gun (Ethicon Endo-Surgery). The sham surgery involved the application of gentle pressure on the stomach with blunt forceps. During the first 3 days of the postoperative period, the animals were fed a liquid diet (Osmolite 1.0 Cal, Abbott Nutrition) and then were returned to the 60% HFD. BW was monitored for 8 weeks after surgery. Body composition was measured using an EchoMRI (Echo Medical Systems) before and 8 weeks postoperatively. Five to 6 weeks after surgery, we performed an OGTT following a 5-6 hour fast (2 g/Kg of 50% dextrose). Female Lgr5^eGFP^ mice were maintained on 60% HFD for 16 weeks to induce obesity, were matched for body weight, and then received sham or VSG surgery as previously described. BW was monitored for 4 weeks after surgery, and the body composition was measured before and 4 weeks after surgery. An OGTT was performed 4 weeks after surgery. Male or female C57BL/6J and Gcg^Tomato^ (GcgCreERT2 mice crossed with Rosa26-tdTomato mice) were utilized to generate mouse enteroids (see below for enteroid methods).

All animal experiments were performed according to an approved protocol by the Institutional Animal Care and Use Committee (IACUC) at the University of Michigan and we followed protocols outlined in the National Institutes of health (NIH) guide for the care and use of laboratory animals (NIH Publications No. 8023, revised 1978).

### Gut peptide assay

At 8 weeks after surgery, control mice were fasted for 5-6 hours and were orally administered a liquid mixed meal (200 µl Ensure plus® supplemented with 30 mg dextrose). At 15 min later, mice were euthanized with CO_2_ gas. The chest cavity was immediately opened, and blood was collected from the left ventricle by the cardiac puncture using EDTA-coated tubes containing DPP-4 inhibitor and aprotinin. Total GLP-1 (MesoScale Discovery, Rockville, MD, USD, cat# K150JVC) was assayed using a sandwich ELISA assay kit, whereas total PYY (Crystal Chem, Elk Grove Village, IL, USA, cat# 81501) were assayed using a standard ELISA assay kit. All assays were performed according to the manufacturer’s instructions.

### Immunohistochemistry (IHC)

The small intestine (duodenum, jejunum, and ileum) from sham and VSG control mice were paraffin-embedded and sectioned onto slides by the University of Michigan In-Vivo Animal Core (IVAC). Paraffin was removed using Citrisolv (VWR). The slides were incubated overnight at 4°C with primary antibody (see above). The slides were stained with corresponding secondary antibodies conjugated to fluorescence dyes for 2 hours at the RT. The slides were mounted in an anti-fade fluorescence mounting medium containing DAPI (Vectashield with DAPI, Vector Laboratories, cat# H-1200). Fluorescent images were obtained using an Olympus IX73 inverted fluorescence microscopy system (Olympus) or Nikon Eclipse Ti fluorescence microscopy with a motorized X-Y stage system. Images were analyzed using Olympus cellSens imaging software (Olympus) or Image Pro 10 software (Media Cybernetics, Rockville, MD, USA).

### FACS

Four-weeks after surgery, a liquid mixed meal was orally administered to overnight fasted Lgr5^eGFP^ mice. At 15 min later, mice were euthanized with CO_2_ gas. The chest cavity was immediately opened, and saline was injected into the left ventricle to perfuse tissues. The small intestine was collected and flushed with cold PBS to remove luminal contents. Each 6 cm region (duodenum, jejunum, and ileum) was opened and either placed into individual tubes containing cold DMEM (high glucose, 4.5 g/l) before immediately being processed for cell dissociation (Stemcell Technologies, Vancouver, BC, Canada, cat# 07174). Dissociated intestinal epithelial cells were stained with antibodies for CD31, CD45, and Annexin V, and the Sytox Blue, and these cells were pre-sorted with MoFlo Astrios cell sorter (Beckman Coulter, Brea, CA, USA) to rule out the immune-, endothelial-, apoptotic-, or dead cells, respectively. The cells were sorted into Norgen lysis buffer (Norgen Biotek, Thorold, ON, Canada #37500) on ice. The cells were vortexed and flash-frozen and stored at -80°C until being used. Intestinal epithelial cells from the WT mice were isolated and stained with APC, Sytox, and GFP for control.

### RNA Library Preparation and Sequencing

Total RNA was isolated from Lgr5^+^ cells, which were sorted by GFP expression, were using the Total RNA Purification Kit (Norgen Biotek). RNA was quantified with the Nanodrop 2000 (ThermoFisher), and RNA integrity was assessed by the Agilent 4200 Tapestation (Agilent Technologies). Libraries were prepared using NEBNext Ultra II Directional Library Prep Kit (New England Biolabs, Ipswich, MA, USA). Sequencing was performed on the NextSeq500 platform (Illumina) at the Biotechnology Research Center at the Cornell University (Ithaca, NY, USA).

### RNA-seq analysis

The sequencing reads were aligned to the mouse genome (mm10). RNA-sequencing reads were mapped to genome release using STAR (v2.5.3a)^40^, and transcript quantification was performed using Salmon (v0.6.0)^41^. Normalization and differential expression analysis were performed using DESeq2^42^. The Benjamini-Hochberg method was used for multiple testing corrections. To define genes altered by VSG vs. sham, samples from each of the regional segments (duodenum, jejunum, and ileum) were analyzed separately. One of the three jejunal samples with sham surgery and one of the three duodenal samples with sham surgery were excluded from the differential expression analysis due to low mapping percentages (below 25%). Two of the four VSG ileal samples and one of the three sham ileal samples were excluded due to low mapping percentages (below 15%). Raw sequencing data and normalized count files are available through GEO accession.

### Mouse enteroid culture

Twelve to 20-week old male or female C57BL/6J or Gcg^Tomato^ mice were euthanized using CO_2_ gas. Duodenal or jejunal segments (∼6 cm) were isolated and placed into falcon tubes containing cold DMEM (high glucose, 4.5 g/l) containing Primocin (Invivogen, cat# ant-pm-1). Intestinal epithelial cells were dissociated using a Gentle Cell Dissociation Reagent (Stemcell Technologies) and were filtered through a 70 µm cell strainer to remove villi. The remaining crypts were resuspended in Matrigel (growth factor reduced) and seeded at the pre-warmed 24-well plate (1000 crypts/20 µl Matrigel). The enteroids were cultured using the IntestiCult™ Organoid Growth Medium for mice and placed in a CO_2_ incubator maintaining 5% CO_2_ concentration and 37°C temperature. The media was changed every 2-3 days, and the enteroids were passaged every 7 days.

At least 7 days after the initial enteroid culture, enteroids were seeded into a 96-well clear-bottom black plate (10-20 enteroids/4-5ul Matrigel/well). After 12-hours of acclimation time, we administered enteroids with chyme, sera, or various bile acids. The chyme and sera were obtained from sham or VSG-treated rats^23^. The chyme from each sham or VSG rats (n=3/surgery) was pooled and filtered with PBS after homogenization (25% w/w)^43^ and administered to mouse duodenal enteroids at 0.1% of vehicle (vol/vol). The sera from each sham or VSG rats (n=3/surgery) was also pooled and administered to mouse duodenal enteroids at 20% of the vehicle (vol/vol). Each bile acid was prepared in DMSO and was administered to mouse duodenal or jejunal enteroids at each dose. For enteroids derived from the C57BL/6J mice, we fixed the enteroids at 12 h, 24 h, or 48 h after treatment using the 4% paraformaldehyde (PFA) solution. The enteroids were blocked with 10% normal donkey serum (NDS) containing PBS and stained overnight with anti-GLP-1 antibody, and then with the secondary antibodies conjugated with fluorescence dyes (FITC or RFP), in consecutive order. For the enteroids derived from the Gcg^Tomato^ mice, we treated 4-hydroxytamoxifen Insolution™ (0.2 µg/ml) along with bile acids. By doing so, the tamoxifen-dependent Cre driver is activated and we could visualize Gcg^+^ cells. The enteroids were imaged using an Olympus IX73 inverted fluorescence microscope system and the images were analyzed using the Olympus cellSense software (Olympus).

### Statistics

GraphPad Prism 8.0 (GraphPad Software, San Diego, CA, USA) was used for the statistical analysis. Data were considered significant when P < 0.05. Where applicable, a two-tailed *t*-test, an ordinary two-way analysis of variance (ANOVA), and a repeated-measures (RM) ANOVA were applied to determine significant main effects and interactions between variables. Significant interactions are indicated in the figure legends, and between-group differences were determined by Tukey’s post hoc testing. A simple linear regression model was used to compare the plasma gut peptide levels by the corresponding EEC number. Data are presented as mean ± SEM. All the padj and p values presented in the RNAseq analysis were determined by the Wald test (DESeq2).

## Acknowledgments

The authors thank Dr. Linda Samuelson (University of Michigan) for providing us with critical opinion on our study. The authors also thank the surgeons for conducting mouse VSG (Alfor Lewis, Andriy Myronovych, Mouhamadoul Toure, and Diana Farris) and Kelli Rule for the technical assistance in mouse breeding, and the In-Vivo Animal Core (IVAC) at UM for tissue processing. This work is supported in part by NIH Awards, DK 121995 (DAS), DK107282 (DAS), DK107652 (RJS), P01-DK117821 (RJS), F32-DK-120159 (BCEP), T32-DK-101357 (BCEP), and T32HD57854-10 (KKL); ADA awards 1-19-IBS-252 (DAS), 1-16-ACE-47 (PS); the Empire State Stem Cell Fund Fellowship (YHH); and by an Early Career Award from the Obesity Society (KSK).

## Author contributions

KSK and DAS conceptualized the study and were responsible for the design of experiments. KSK, BCEP, YHH, KKL, PHD, and LW were responsible for executing the experiments. KSK, LCS, JRS, PS, and DAS were responsible for analysis and interpretation of data. KSK and DAS were responsible for drafting of the manuscript. DAS provided final approval of the submitted manuscript.

## References

1. Estimate of Bariatric Surgery Numbers, 2011-2017 | American Society for Metabolic and Bariatric Surgery.

2. Schauer, P. R. et al. Bariatric Surgery versus Intensive Medical Therapy in Obese Patients with Diabetes. N. Engl. J. Med. 366, 1567–1576 (2012).

3. Kim, K.-S. & Sandoval, D. A. Endocrine function after bariatric surgery. in Comprehensive Physiology vol. 7 783–798 (John Wiley & Sons, Inc., 2017).

4. Peterli, R. et al. Metabolic and hormonal changes after laparoscopic Roux-en-Y gastric bypass and sleeve gastrectomy: A randomized, prospective trial. Obes. Surg. 22, 740–748 (2012).

5. Dimitriadis, E. et al. Alterations in gut hormones after laparoscopic sleeve gastrectomy: A prospective clinical and laboratory investigational study. Ann. Surg. 257, 647–654 (2013).

6. Nannipieri, M. et al. Roux-en-Y gastric bypass and sleeve gastrectomy: mechanisms of diabetes remission and role of gut hormones. J. Clin. Endocrinol. Metab. 98, 4391–4399 (2013).

7. Grayson, B. E., Schneider, K. M., Woods, S. C. & Seeley, R. J. Improved rodent maternal metabolism but reduced intrauterine growth after vertical sleeve gastrectomy. Sci. Transl. Med. 5, 199ra112 (2013).

8. Schauer, P. R. et al. Bariatric Surgery versus Intensive Medical Therapy for Diabetes — 5-Year Outcomes. N. Engl. J. Med. 376, 641–651 (2017).

9. Cavin, J.-B. et al. Differences in Alimentary Glucose Absorption and Intestinal Disposal of Blood Glucose After Roux-en-Y Gastric Bypass vs Sleeve Gastrectomy. Gastroenterology 150, 454–64.e9 (2016).

10. Nausheen, S., Shah, I. H., Pezeshki, A., Sigalet, D. L. & Chelikani, P. K. Effects of sleeve gastrectomy and ileal transposition, alone and in combination, on food intake, body weight, gut hormones, and glucose metabolism in rats. Am. J. Physiol. Endocrinol. Metab. 305, E507–18 (2013).

11. Li, F., Peng, Y., Zhang, M., Yang, P. & Qu, S. Sleeve gastrectomy activates the GLP-1 pathway in pancreatic β cells and promotes GLP-1-expressing cells differentiation in the intestinal tract. Mol. Cell. Endocrinol. 436, 33–40 (2016).

12. Demitrack, E. S. & Samuelson, L. C. Notch regulation of gastrointestinal stem cells. J. Physiol. 594, 4791–4803 (2016).

13. VanDussen, K. L. et al. Notch signaling modulates proliferation and differentiation of intestinal crypt base columnar stem cells. Development 139, 488–497 (2012).

14. Ryan, K. K. et al. FXR is a molecular target for the effects of vertical sleeve gastrectomy. Nature 509, 183–188 (2014).

15. Myronovych, A. et al. Vertical sleeve gastrectomy reduces hepatic steatosis while increasing serum bile acids in a weight-loss-independent manner. Obesity 22, 390–400 (2014).

16. Nakatani, H. et al. Serum bile acid along with plasma incretins and serum high–molecular weight adiponectin levels are increased after bariatric surgery. Metabolism 58, 1400–1407 (2009).

17. Dossa, A. Y. et al. Bile acids regulate intestinal cell proliferation by modulating EGFR and FXR signaling. Am. J. Physiol. Gastrointest. Liver Physiol. 310, G81–92 (2016).

18. Fu, T. et al. FXR Regulates Intestinal Cancer Stem Cell Proliferation. Cell 176, 1098–1112.e18 (2019).

19. Kim, K. S. et al. Glycemic effect of pancreatic preproglucagon in mouse sleeve gastrectomy. JCI Insight 4, e129452 (2019).

20. Cao, X., Flock, G., Choi, C., Irwin, D. M. & Drucker, D. J. Aberrant Regulation of Human Intestinal Proglucagon Gene Expression in the NCI-H716 Cell Line. (2003) doi:10.1210/en.2002-0049.

21. Hill, M. E., Asa, S. L. & Drucker, D. J. Essential Requirement for iPax/i 6 in Control of Enteroendocrine Proglucagon Gene Transcription. Mol. Endocrinol. 13, 1474–1486 (1999).

22. Gehart, H. et al. Identification of Enteroendocrine Regulators by Real-Time Single-Cell Differentiation Mapping. Cell 176, 1158–1173.e16 (2019).

23. Arble, D. M. et al. Metabolic comparison of one-anastomosis gastric bypass, single-anastomosis duodenal-switch, Roux-en-Y gastric bypass, and vertical sleeve gastrectomy in rat. Surg. Obes. Relat. Dis. 14, 1857–1867 (2018).

24. Gimeno, R. E., Briere, D. A. & Seeley, R. J. Leveraging the Gut to Treat Metabolic Disease. Cell Metabolism vol. 31 679–698 (2020).

25. Chambers, A. P. et al. Regulation of gastric emptying rate and its role in nutrient-induced GLP-1 secretion in rats after vertical sleeve gastrectomy. AJP Endocrinol. Metab. 306, E424–E432 (2014).

26. Steinert, R. E. et al. Ghrelin, CCK, GLP-1, and PYY(3–36): secretory controls and physiological roles in eating and glycemia in health, obesity, and after RYGB. Physiol. Rev. 97, 411–463 (2017).

27. Wölnerhanssen, B. K. et al. Deregulation of transcription factors controlling intestinal epithelial cell differentiation; A predisposing factor for reduced enteroendocrine cell number in morbidly obese individuals. Sci. Rep. 7, (2017).

28. Mumphrey, M. B., Hao, Z., Townsend, R. L., Patterson, L. M. & Berthoud, H.-R. Sleeve Gastrectomy Does Not Cause Hypertrophy and Reprogramming of Intestinal Glucose Metabolism in Rats. Obes. Surg. 25, 1468–1473 (2015).

29. Beyaz, S. et al. High-fat diet enhances stemness and tumorigenicity of intestinal progenitors. Nature 531, 53–58 (2016).

30. Sakar, Y. et al. Impact of high-fat feeding on basic helix-loop-helix transcription factors controlling enteroendocrine cell differentiation. Int. J. Obes. 38, 1440–1448 (2014).

31. Wilson-Pérez, H. E. et al. The effect of vertical sleeve gastrectomy on food choice in rats. Int. J. Obes. (Lond). 37, 288–295 (2013).

32. Shin, A. C. et al. Longitudinal assessment of food intake, fecal energy loss, and energy expenditure after roux-en-y gastric bypass surgery in high-fat-fed obese rats. Obes. Surg. 23, 531– 540 (2013).

33. Carulli, A. J. et al. Notch receptor regulation of intestinal stem cell homeostasis and crypt regeneration. Dev. Biol. 402, 98–108 (2015).

34. Myronovych, A. et al. The role of small heterodimer partner in nonalcoholic fatty liver disease improvement after sleeve gastrectomy in mice. Obesity 22, 2301–2311 (2014).

35. Osinski, C. et al. Type 2 diabetes is associated with impaired jejunal enteroendocrine GLP-1 cell lineage in human obesity. Int. J. Obes. 45, 170–183 (2021).

36. Wilson-Pérez, H. E. et al. Vertical sleeve gastrectomy is effective in two genetic mouse models of glucagon-like peptide-1 receptor deficiency. Diabetes 62, 2380–2385 (2013).

37. Boland, B. et al. The PYY/Y2R-deficient mouse responds normally to high-fat diet and gastric bypass surgery. Nutrients 11, (2019).

38. Behary, P. et al. Combined GLP-1, oxyntomodulin, and peptide YY improves body weight and glycemia in obesity and prediabetes/type 2 diabetes: A randomized, single-blinded, placebo-controlled study. Diabetes Care 42, 1446–1453 (2019).

39. Finan, B. et al. A rationally designed monomeric peptide triagonist corrects obesity and diabetes in rodents. Nat. Med. 21, 27–36 (2014).

40. Dobin, A. et al. STAR: ultrafast universal RNA-seq aligner. Bioinformatics 29, 15–21 (2013).

41. Patro, R., Duggal, G., Love, M. I., Irizarry, R. A. & Kingsford, C. Salmon provides fast and bias-aware quantification of transcript expression. Nat. Methods 14, 417–419 (2017).

42. Love, M. I., Huber, W. & Anders, S. Moderated estimation of fold change and dispersion for RNA-seq data with DESeq2. Genome Biol. 15, 550 (2014).

43. Jørgensen, J. R. & Mortensen, P. B. Influence of feces from patients with ulcerative colitis on butyrate oxidation in rat colonocytes. Dig. Dis. Sci. 44, 2099–2109 (1999).

