## Supplementary materials for "Vertical Sleeve Gastrectomy Induces Enteroendocrine Cell Differentiation of Intestinal Stem Cells Through Farnesoid X Receptor Activation"

**Supplementary Material**

### Supplementary figure legends

**Fig. S1.** Lgr5eGFP mice VSG phenotype. (A) Schematic experimental timeline. (B) Representative immunofluorescence image of the crypt of the Lgr5eGFP mice jejunum (bar = 100 µm). (C-E) BW change (C), fat mass (D), and lean mass (E) in sham or VSG-treated Lgr5eGFP mice. (F) Oral glucose tolerance (2 g/Kg) of sham or VSG mice. Mean ± SEM. Statistics, t-test or ANOVA. **P<0.01, n=8-9.

**Fig. S2.** Lgr5eGFP mice VSG-FACS results. (A) Pre-sorted Sytox- and APC+ intestinal epithelial cells (%) of the sham or VSG-treated Lgr5eGFP mice. (B) EGFP+ intestinal epithelial cells (%) of the sham or VSG-treated Lgr5eGFP mice. Main effect of surgery. * P<0.05, n=5-6.

**Fig. S3.** Lgr5eGFP mice VSG-RNAseq results. (A-B) Venn diagram of up- (A) or downregulated (B) genes that are unique or shared across regional segments with VSG surgery compared to sham (upregulated genes defined by p value < 0.05, adjusted P < 0.2, normalized counts > 50, $\log_2$ fold change > 0.5). n=2-4 samples/surgery type for each regional segments. (C-E) Principal component analysis (PCA) of RNAseq data from duodenal (C), jejunal (D), or ileal (E) $GFP^+$ ISCs under sham and VSG surgery. n=2-4 samples/surgery type for each regional segments. (F) Volcano plot of differentiatlly expressed genes (348 up; 162 down) in jejunal $GFP^+$ ISCs under VSG relative to sham surgery (p value < 0.05, padj < 0.2, $\log_2$ fold change > 0.5, and normalized counts > 50). Dashed lines denote p value and $\log_2$ fold change threshold (n=4 in VSG and n=2 in sham).

**Fig. S4.** qPCR validation of the RNAseq results. Gcg and Pyy gene expression from $GFP^+$ ISCs of the duodenum, jejunum, ileum of sham or VSG Lgr5eGFP mice. Statistics, *t*-test. *P<0.05, n=5-6.

**Fig. S5.** The goblet cell number in the sham vs. VSG mouse intestine. (A-C) Muc2+ cell number of the intestinal transverse sections (sec.) of the duodenum (A), jejunum (B), or ileum (C). n=8.

**Fig. S6.** Cholic acid (CA) treatment increases the number of enteroids containing GLP-1+ EECs in a dose-dependent manner. (A-B) Gcg-tomato mouse enteroids were exposure to the CA for 24 h. Violin plot of the individual enteroids (n=1000 enteroids/well) indicates median and quartiles (dotted lines). Numbers (%) above each violin plot incidate the proportion of the enteroids containing GLP-1+ EECs. Statistics, ANOVA. *P<0.05, ***P<0.001.

**Fig. S7.** Cholic acid (CA) treatment increases the number of enteroids containing GLP-1+ EECs in a GLP-1 receptor dependent manner. (A-C) Mouse enteroids were exposure to the CA alone or with a potent GLP-1 receptor antagonist, exendin 9-39 (Ex9) for 48 h, and the average enteroids area (A), GLP-1+ cell number per enteroids (B), and GLP-1+cell number per enteroid area (GLP-1+ cell density; C) were measured. n=4/group, each well contains 20 enteroids. Mean ± SEM. Statistics, ANOVA. *P<0.05, **P<0.01.

**Fig. S8.** Cholic acid (CA), but not muricholic acid (MCA) treatment increases the GLP-1+ EECs density. (A-B) CA treatment did not change the average enteroid area (A) while it significantly increased GLP-1+ cell density (B). (C-D) MCA treatment dose-dependently increased the average enteroid area (C) while it did not change the GLP-1+ cell density (D). n=3-5/group, each well contains 20 enteroids. Mean ± SEM. Statistics, ANOVA. *P<0.05, **P<0.01, ***P<0.001.

### Supplementary Figures

Supplementary Figure 1.

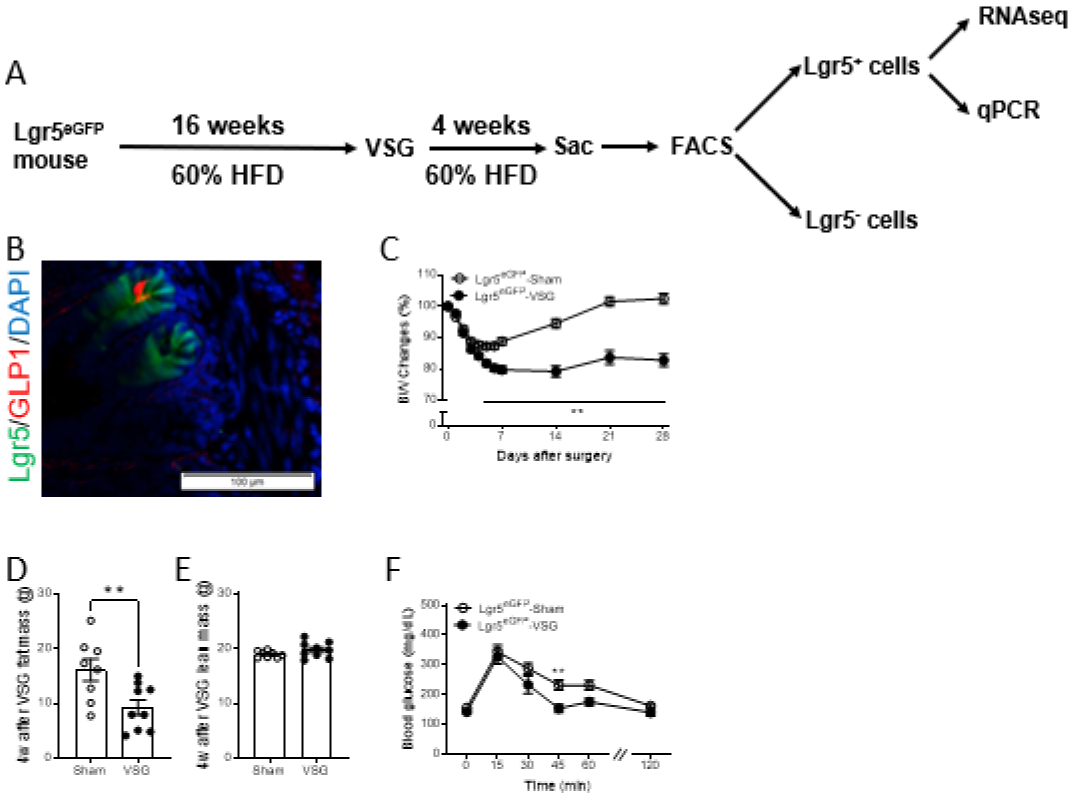

Supplementary Figure 2.

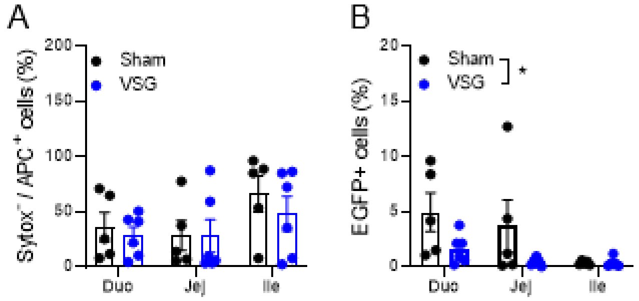

Supplementary Figure 3.

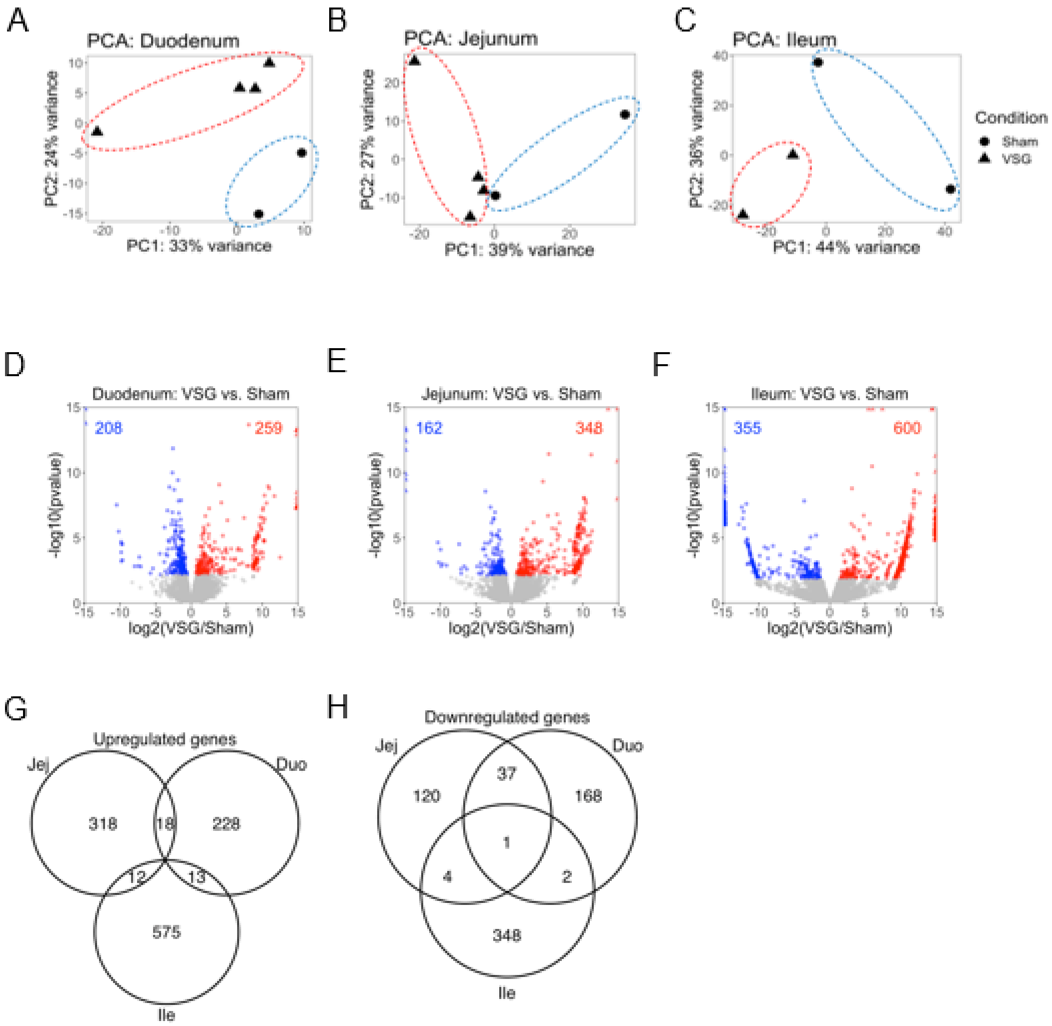

Supplementary Figure 4.

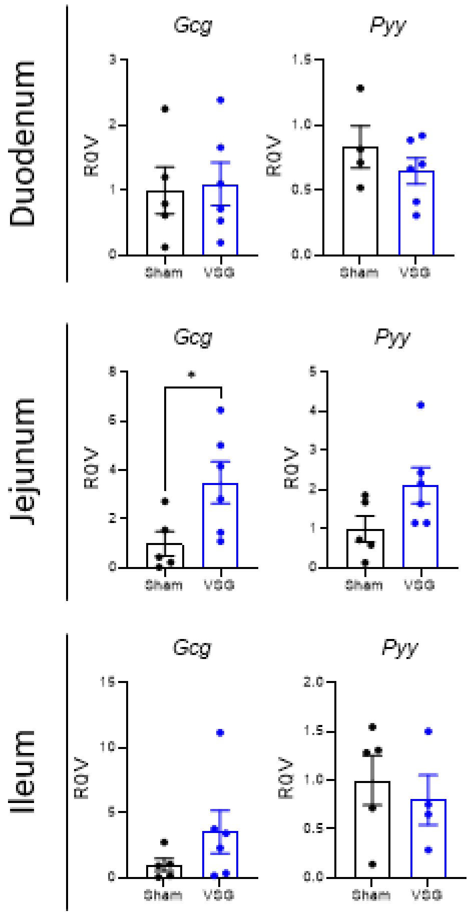

Supplementary Figure 5.

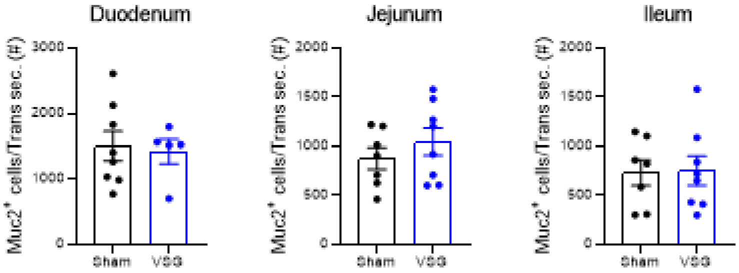

Supplementary Figure 6.

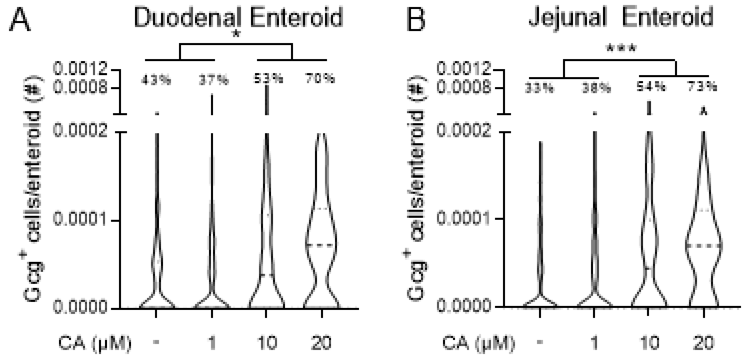

Supplementary Figure 7.

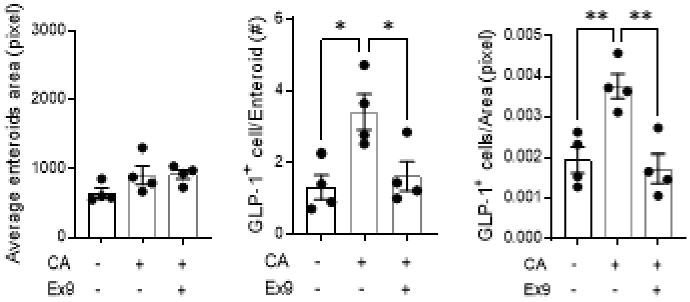

Supplementary Figure 8.

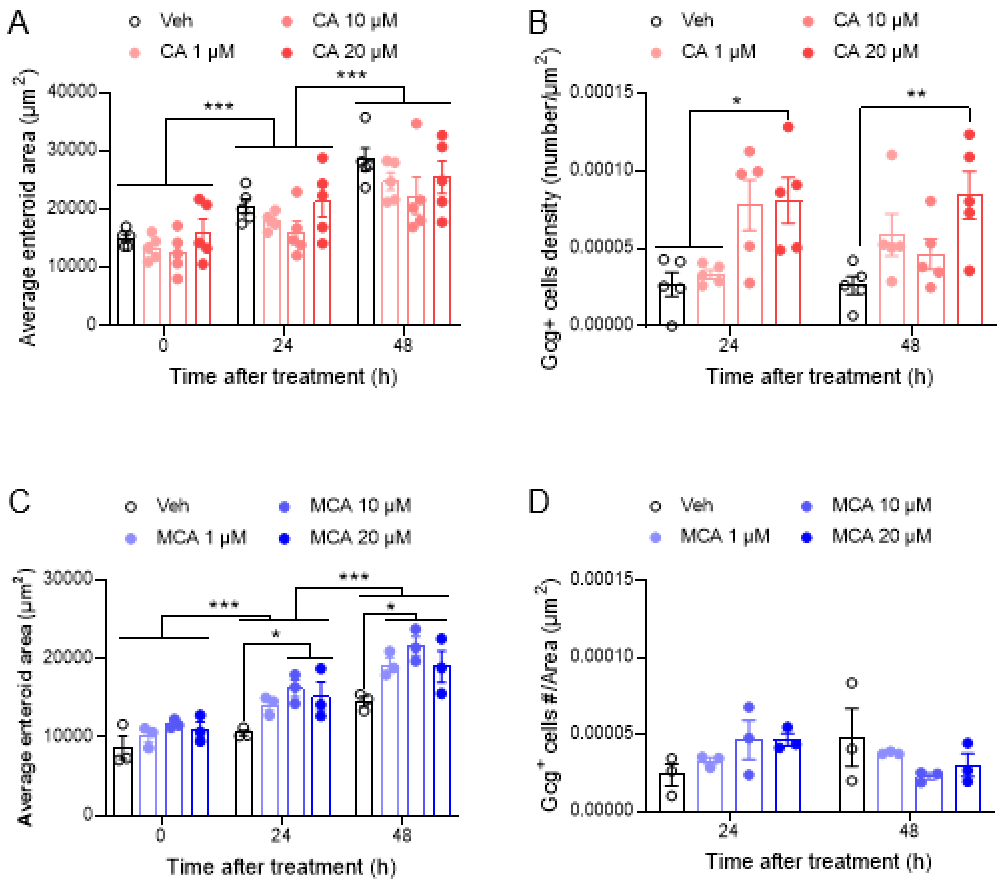
